# Microbial communities from weathered outcrops of a sulfide-rich ultramafic intrusion, and implications for mine waste management

**DOI:** 10.1101/2022.10.03.510692

**Authors:** Kathryn K. Hobart, ZhaaZhaawaanong Greensky, Kimberly Hernandez, Joshua M. Feinberg, Jake V. Bailey, Daniel S. Jones

**Affiliations:** Dept. Of Earth & Environmental Sciences, University of Minnesota, 116 Church Street SE, Minneapolis, MN 55455, U.S.A.; Institute for Rock Magnetism, University of Minnesota, 116 Church Street SE, Minneapolis, MN 55455, U.S.A.; Dept. of Earth and Environmental Science, New Mexico Institute of Mining and Technology, 314 Mineral Science and Engineering Complex, Socorro, NM 87801, U.S.A; National Cave and Karst Research Institute, 400-1 Cascades Avenue, Carlsbad, NM, 88220, U.S.A.

**Keywords:** Duluth Complex, sulfide mineral, pyrrhotite, *Sulfuriferula*, weathering

## Abstract

The Duluth Complex, Northeastern Minnesota, contains sulfide-rich magmatic intrusions that, collectively, represent one of the world’s largest known economic deposits of copper, nickel, and platinum group elements (Cu-Ni-PGEs). Previous work showed that microbial communities associated with experimentally-weathered Duluth Complex waste rock and tailings were dominated by uncultivated taxa and other populations not typically associated with mine waste. However, those experiments were designed for kinetic testing and do not necessarily represent the conditions expected for reclaimed mine waste or long-term weathering in the environment. We therefore used 16S rRNA gene methods to characterize the microbial communities present on the surfaces of naturally-weathered and historically disturbed outcrops of Duluth Complex material, as well as a circumneutral seep draining a reclaimed blast pit. Rock surfaces were dominated by diverse uncultured *Ktedonobacteria, Acetobacteria*, and *Actinobacteria* while seeps were dominated by *Proteobacteria*, including *Leptothrix* spp. and *Methylovulum* spp. All samples had abundant algae and other phototrophs. These communities were distinct from previously-described microbial assemblages from experimentally-weathered Duluth Complex rocks, suggested different energy and nutrient resources in the reclaimed rocks, outcrops, and seeps. Sulfide mineral incubations performed with and without algae showed that photosynthetic microorganisms could have an inhibitory effect on some of the autotrophic populations from the site, resulting in slightly lower sulfate release and differences in the dominant microorganisms. The microbial assemblages from these weathered outcrops show how communities are expected to develop during natural weathering of sulfide-rich Duluth Complex rocks, and represent baseline data that could be used to evaluate the effectiveness of future reclamation of tailings and waste rock produced by large scale mining operations.

## 1. INTRODUCTION

Microorganisms are important catalysts for sulfide mineral oxidation and dissolution in natural and engineered settings. Their role in generating acid rock drainage is frequently taken into account when examining the risks of proposed, current, and legacy mining activities (Schippers *et al*., 1996; Edwards *et al*., 1999; Schippers and Sand, 1999; Baker and Banfield, 2003; Ňancucheo and Johnson, 2011). Under extremely acidic conditions (pH < 4), iron- and sulfur-oxidizing microorganisms regenerate the oxidant Fe(III) and produce acids that accelerate the oxidation of metal sulfide minerals and intensify the production of acidic and metal-rich waste streams (Schippers *et al*., 1996; Schippers and Sand, 1999; Nordstrom *et al*., 2015). Understanding the importance of microbial processes in acidic, sulfide mineral-impacted systems has led to new strategies for mineral extraction, mine waste remediation, and prevention of acid rock drainage (Dugan and Apel, 1983; Onysko *et al*., 1984; Schippers and Sand, 1999; Rohwerder *et al*., 2003; Rawlings and Johnson, 2007; Brune and Bayer, 2012; Vera *et al*., 2013; Johnson, 2018).

The Duluth Complex, located in Northern Minnesota, USA, is a layered mafic intrusion that hosts some of the largest undeveloped Cu-Ni-PGE deposits in the world (Thériault *et al*., 2000; Miller *et al*., 2002; Severson *et al*., 2002). The relatively low total sulfide mineral content of these deposits and the buffering capacity of the host silicate minerals means that weathering of these rocks is not expected to produce extremely acidic conditions (Lapakko, 1988, 2015; Seal *et al*., 2015). Field and laboratory testing of experimentally-generated tailings and waste rock produces moderately acidic leachate (pH 4 to 7), only rarely reaching pH values below pH 4 (Lapakko and Antonson, 2012). Unlike highly acidic systems, the microbial role in sulfide mineral oxidation under mildly acidic to circumneutral conditions is not well understood, and until recently (Percak-Dennett *et al*., 2017; Napieralski *et al*., 2022), microorganisms were not thought to significantly accelerate the oxidation of the acid-insoluble sulfide pyrite (FeS_2_) above pH 4 (Arkesteyn, 1980; Nordstrom, 1982; Schippers and Jørgensen, 2002; Schippers, 2004; Korehi *et al*., 2014). The microbial communities found in circumneutral sulfidic mine waste are not as well understood as those found in extremely acidic environments, and are often characterized by the abundance of sulfur-oxidizing rather than iron-oxidizing bacteria (Schippers *et al*., 1996, 1996; Lindsay *et al*., 2009; Chen *et al*., 2013), and frequently contain uncultivated taxa (Mendez *et al*., 2008; Chen *et al*., 2013; Korehi *et al*., 2014). Furthermore, the most abundant primary sulfide mineral in Duluth Complex deposits is pyrrhotite (Fe_1-x_S, where 0 ≤ x ≤ 0.125), for which biological oxidation rates have not been established as extensively as for pyrite, especially at high pH (e.g., Belzile *et al*., 2004).

Recent work showed that long-term laboratory and field-leaching experiments with waste rock and tailings from Duluth Complex materials contained diverse microbial communities that were populated by organisms not typically associated with mine waste (Jones, Lapakko, *et al*., 2017). These communities experiments included numerous 16S rRNA gene and transcript sequences from groups that are only known from environmental samples. Further, the microbial communities sampled from these experiments were primarily composed of taxa associated with organoheterotrophic or sulfur-oxidizing lifestyles and had a conspicuous absence of known iron-oxidizing taxa that are typically implicated in sulfide mineral dissolution.

However, these weathering experiments were performed in a controlled setting that is appropriate for kinetic testing but might not necessarily mimic the conditions encountered in large-scale tailings and waste rock piles or leach pads (Lapakko, 2015; Maest and Nordstrom, 2017). Furthermore, once mine waste is stabilized by planting or reclaimed following mine closure, sulfide mineral-associated microbial communities are subjected to very different redox conditions and nutrient inputs through plant colonization and soil development. Therefore, we characterized microbial communities associated with naturally weathered sulfide-bearing Duluth Complex rocks that are exposed in outcrops in Northern Minnesota, including from a former blast pit where rocks were extracted in 1974 and has since been reclaimed. We compare the microbial communities that developed on rock surfaces and in seeps and associated waters from these areas to communities described in earlier field and laboratory weathering experiments. We also observed abundant algae at the sites. In light of previous studies that show that organic inputs by algae can limit sulfide mineral oxidation (Das *et al*., 2009; Johnson, 2014; Bwapwa *et al*., 2017; Gruzdev *et al*., 2020; Rambabu *et al*., 2020), we compared how algal growth affected sulfide mineral oxidation in laboratory incubations with communities from the site.

## 2. RESULTS

### 2.1 Samples, field observations, and geochemistry

Weathered rock, sediment, and water samples were collected from four sites in Northern Minnesota. All sites are located in the South Kawishiwi Intrusion of the Duluth Complex (Severson, 1994). The “INCO Pit” site (IB samples) is a surface outcrop of ore-bearing Duluth Complex material. In 1974, the International Nickel Company (INCO) removed roughly 10,000 tons of ore-grade material as part of an application to mine in this area; the site was reclaimed when the application was withdrawn in 1975. We collected water and sediment from a seep flowing out of the reclaimed material, and water and algal biomass from a stream approximately 10m from the seep. Effluent from the seep was pH 6.42 at the time of sampling. We also collected exposed, weathered rock from the northeast edges of the reclaimed bulk sample site. Samples collected from this area also include naturally-weathered Duluth Complex material approximately a quarter kilometer from the reclaimed test pit, at the “Gravel Pit” site (GG samples). Here, glacial till deposits were extracted for gravel as part of road-building operations in the 1940s, exposing the glaciated Duluth Complex surface to weathering. We sampled the exposed, crumbly weathered rock at this site, as well as water and biomass from algae-filled drill holes in exposed but consolidated weathered rock. Samples are summarized in Table 1, and concentration of dissolved anions from the three aqueous samples collected at this site are reported in Table S1 in the supplementary information.

**Table 1.** Samples collected for this study

| Name | Site | Description |
| --- | --- | --- |
| IB-2 | INCO Pit | Sediment from a pH 6.42 seep |
| IB-4 | INCO Pit | Algal biomass from stream below reclaimed test pit area |
| IB-5 | INCO Pit | Weathered, poorly-consolidated Duluth Complex rock |
| IB-6 | INCO Pit | Weathered, poorly-consolidated Duluth Complex rock |
| GG-1 | Gravel Pit | Weathered, poorly-consolidated Duluth Complex rock |
| GG-2 | Gravel Pit | Water and algal biomass from inside a drill hole in weathered (but consolidated) Duluth Complex rock |
| HB-1 | Hammer Breaker | Weathered Duluth Complex rock |
| HB-2 | Hammer Breaker | Weathered Duluth Complex rock |
| DO-1 | Duluth Outcrop | Weathered Duluth Complex rock |
| DO-2 | Duluth Outcrop | Weathered Duluth Complex rock |
| DO-3 | Duluth Outcrop | Weathered Duluth Complex rock |
| DO-4 | Duluth Outcrop | Weathered Duluth Complex rock |

We also collected samples of weathered Duluth Complex material from the surfaces of two additional roadside outcrops: the “Hammer Breaker” outcrop of the Duluth Complex along Forest Rte. 429 (HB samples), and from an outcrop of Duluth Complex material off St. Louis Co. Road 623 (DO samples), which was exposed when the road was re-routed to allow for the expansion of the Cliffs Natural Resources Northshore mine. Samples from this site are listed in Table 1. See Figure 1 for field images of sampling sites, and Figure S1 and S2 in the supplementary information for spatial relationship between sampling sites, geographic locations, and additional field images.

**Figure 1.**
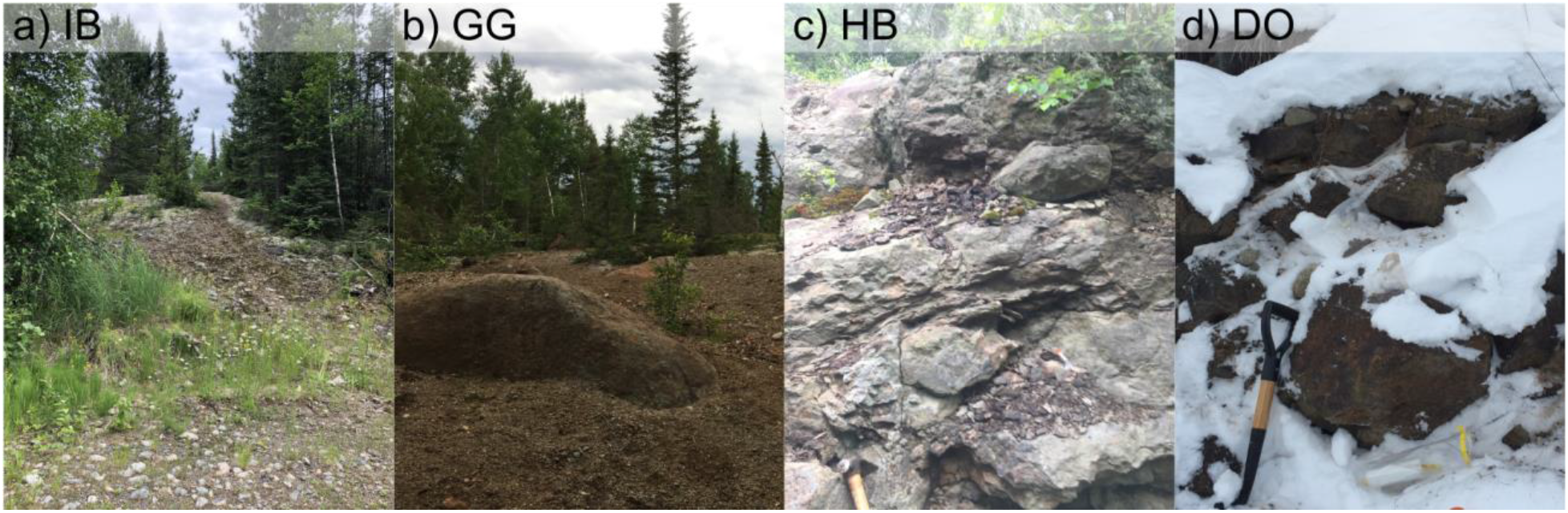
Field images of the (a) IB, (b) GG, (c) HB, and (d) DO sampling sites.

### 2.2 16S rRNA gene libraries from field samples

We generated 20 rRNA gene amplicon libraries from 12 samples of weathered, sulfide-bearing Duluth Complex material collected from four sites (Table 1). The libraries had between 18,504 and 158,184 sequences per sample, with an average library size of 52,703 sequences (standard deviation 37,929). Replicate libraries produced with 25 and 30 PCR cycles for the first amplification step were similar (Figure S3), so we only report results from the 25-cycle libraries. Libraries are available in the Sequence Read Archive (https://www.ncbi.nlm.nih.gov/sra) under accession PRJNA885421.

Several operational taxonomic units (OTUs) were abundant across libraries, with chloroplasts representing many of the most abundant OTUs. To evaluate how the presence of algae and other eukaryotes affected sample clustering and to more closely investigate the non-photosynthetic microbial community, hierarchical agglomerative cluster analysis was performed with and without OTUs identified as chloroplasts, mitochondria, eukaryotes, and an abundant *Gammaproteobacteria* OTU that represents a protist symbiont (OTU 2, discussed below) (Fig. 2, Fig. S4).

**Figure 2.**
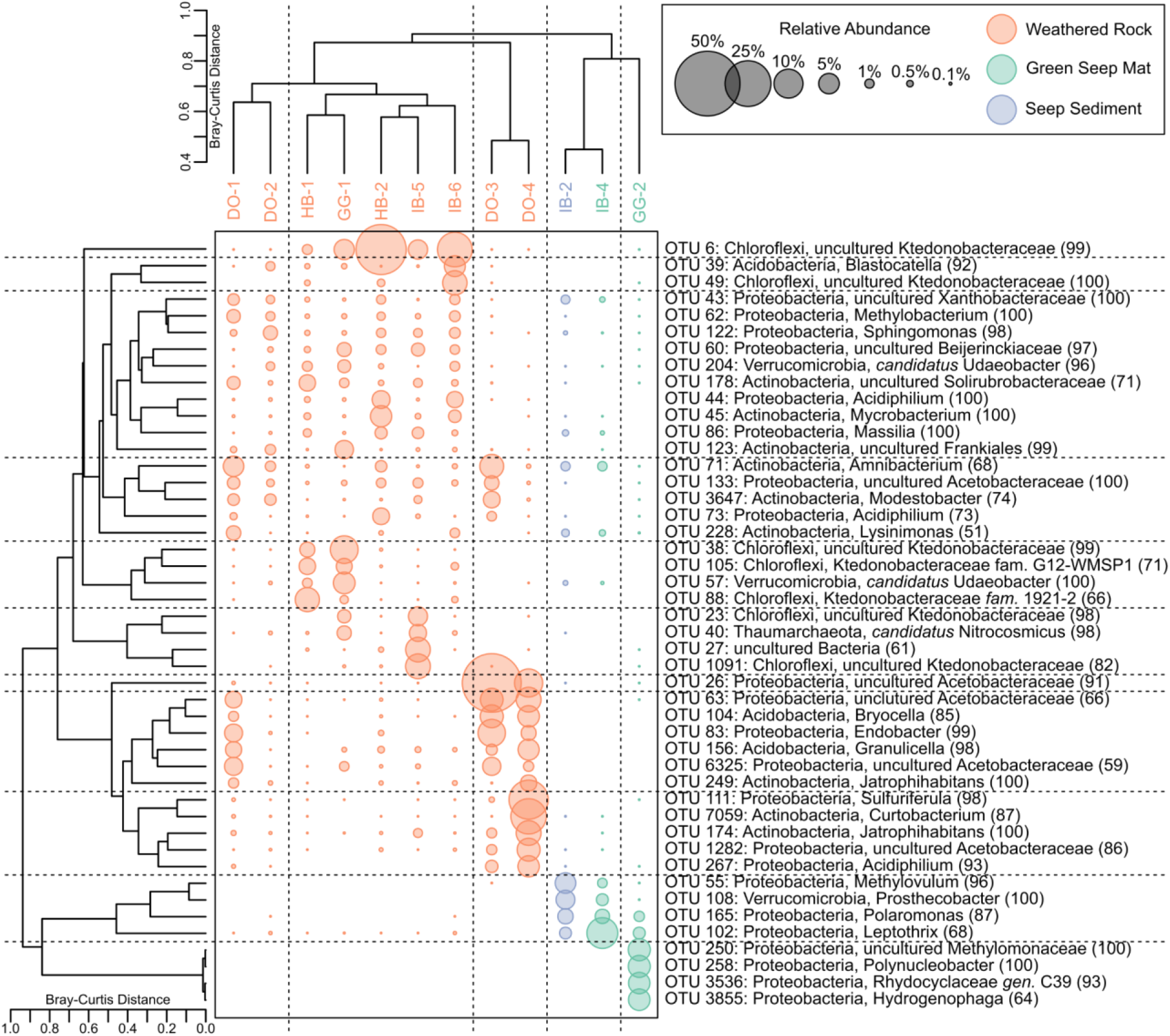
Hierarchical agglomerative cluster analyses of libraries collected for this study, with OTUs identified as chloroplasts, mitochondria, and eukaryotes removed. Sizes of the points scales with the relative abundance of the OTUs. The Q-mode cluster analysis was calculated with all OTUs, while the R-mode cluster analysis only included OTUs abundant in at least one sample at 5% or greater abundance. The taxonomic affiliation of each OTU includes its phylum- and genus-level classification, with confidence scores provided in parentheses. OTUs that are unclassified at the genus level are identified with the highest available taxonomic classification. An equivalent analysis including all OTUs is presented in Figure S4.

Without chloroplasts and other eukaryote-affiliated OTUs (Fig. 2), all weathered rock samples clustered together. The seep sediment and green mat samples from the INCO reclaimed pit site (IB-2 and IB-4) cluster separately from the weathered rock libraries (Fig. 2). Similarly, GG-2, the library generated from the algal biomass collected from drill holes in the glaciated surface site (Fig. 1b, Fig. S2), clusters separately from other libraries, and is more similar to the seep sediment and green seep mat libraries than the weathered rock libraries. The seep sediment and algal biomass samples from both the IB and GG sites contain unique non-chloroplast OTUs in addition to small numbers of the OTUs that are most abundant in the weathered rock samples. When chloroplasts, mitochondria, and eukaryotes are included (Fig. S4), the weathered rock samples continue to cluster separately from the seep sediment and algal mat samples, indicating that these samples are separated by distinct groups of both photosynthetic and non-photosynthetic taxa.

IB-2 and IB-4 share two abundant OTUs (OTU 95, chloroplast, and OTU 102, *Leptothrix* spp.) that separate these samples from the weathered rock samples taken from the same site (IB-5 and IB-6, Fig. 2). Similarly, GG-2, the library generated from the algal biomass collected from the drill hole at glaciated DC outcrop (Fig. 1b), clusters separately from weathered rock collected at the same site (GG-1). GG-2 is dominated by an OTU (OTU 5) that is most closely related (99.2% similarity) to a chloroplast from *Euglena granulata* (strain UTEX 2345, Kosmala *et al*., 2009), and an OTU (OTU 2) initially classified only as a gammaproteobacterium. By placing OTU 2 into an ARB (Ludwig *et al*., 2004) database (SILVA version 138.1, Quast *et al*., 2013) showed that this OTU is a member of an “*incertae sedis*” group in the *Gammaproteobacteria* and clusters in a sister clade to *candidatus* Ovatusbacter (Dirren and Posch, 2016). A similar environmental sequence, as determined by BLASTN search of the nr/nt National Center for Biotechnology Information database (NCBI Resource Coordinators, 2016), is associated with the euglenid algae *Trachelomonas scabra*. This suggests OTU 2 is a proteobacterial symbiont of the euglenid algae in the green mat. Similar gammaproteobacterial symbionts of *Trachelomonas scabra* have been associated with euglenid biofilms in ARD environments (e.g., Jones *et al*., 2015). Because the abundance of OTU 2 matches the abundance of the euglenid algae in this sample (OTU 5), and given its likely role as a symbiont of that algae, we removed OTU 2 along with the chloroplast, mitochondria, and eukaryote OTUs in the multivariate statistical analyses shown in Figures 2 and 3.

**Figure 3.**
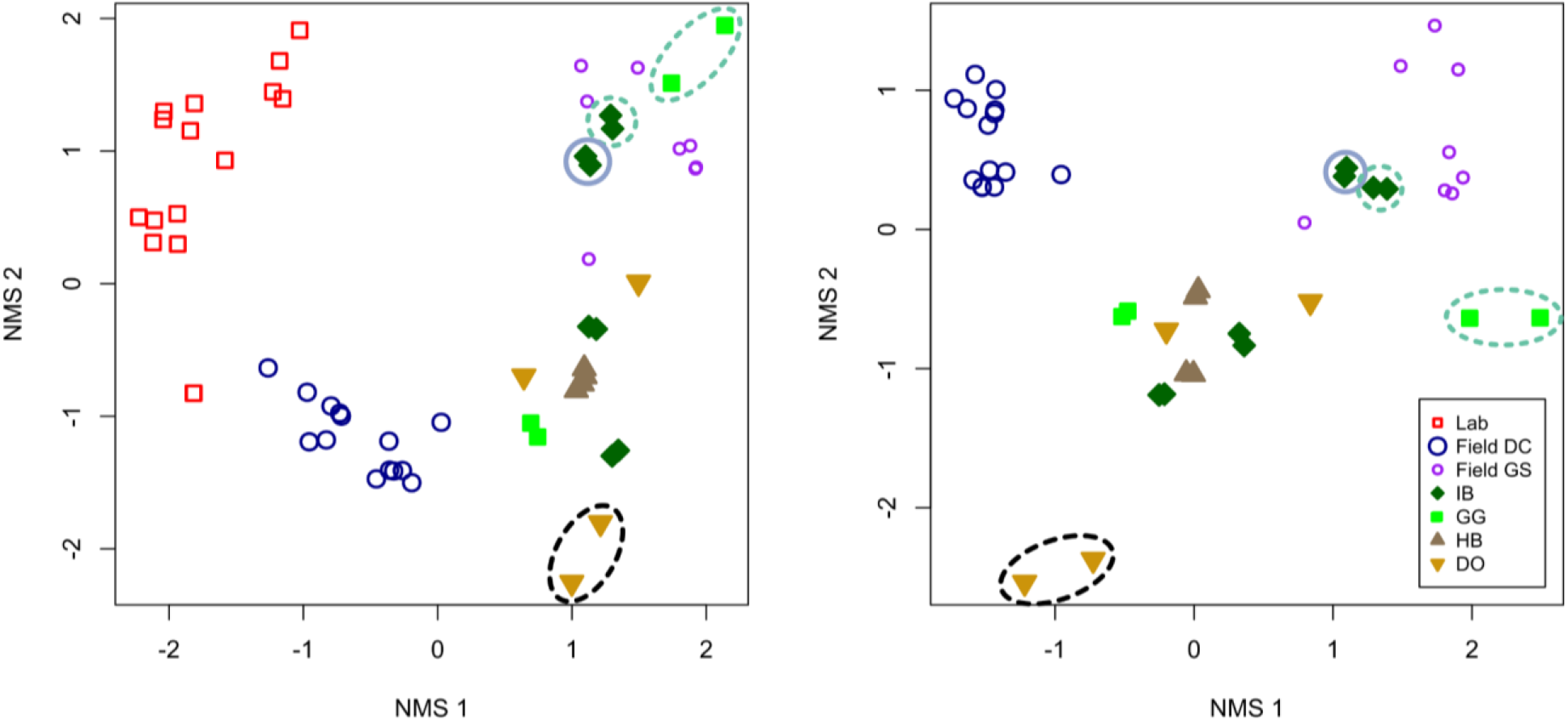
(a) NMS ordinations of rRNA amplicon libraries from naturally-weathered Duluth Complex outcrops (libraries IB, GG, HB, and DO, this study), laboratory humidity cells, and experimental field rock piles, with OTUs identified as chloroplasts, mitochondria, and eukaryotes removed. Libraries from laboratory experiments (“lab,” red open squares), and experimental field rock piles (“Field DC,” Duluth Complex material, dark blue larger circles, and “Field GS,” Ely Greenstone material, purple smaller circles) are from (Jones, Lapakko, *et al*., 2017). (b) NMS ordinations of rRNA amplicon libraries from the experimental field rock piles and naturally weathered outcrops only. Stress for (a) is 7.5, stress for (b) is 5.8. Sky blue solid circled points indicate seep sediment samples (IB-2), teal green dotted circled points indicate water and algal biomass samples (IB-4, GG-2). All other IB, GG, HB, and DO samples were collected from weathered rock. The black dashed circle indicates DO samples that are separated from the other weathered rock samples in the hierarchical agglomerative cluster analysis (Figure 2).

All other IB, GG, and HB libraries were generated from weathered rock samples, and cluster together (Fig. 2, Fig. S5). In addition to the most abundant chloroplast OTUs in this cluster (OTUs 3, 9, and 52), these samples contained abundant uncultured representatives of the family *Ktedonobacteraceae* (OTUs 6, 38, 49, 88, and 105), genus *Acidiphilum* (OTU 44), and uncultured representatives of the *Solirubrobacteraceae*, (OTU 178), *Beijerinckiaceae* (OTU 60), and *Verrucomicrobia* (*candidatus* Udaeobacter, OTU 57) (Fig. 2).

The libraries generated from weathered rock samples DO-3 and DO-4 cluster separately from the IB, GG, and HB samples (Fig. 2). These libraries contain abundant OTUs that are only minor components of the IB and HB libraries (<1%). Two of the most abundant OTUs in this group (OTU 13 and 70) are most closely related to mitochondrial rRNA genes of the ascomycetous fungi *Leotiomycetes* (92% similar to *Leotiomycetes* sp. A910, Muggia *et al*., 2016) and *Chaetothyriales* (86% similar to *Chaethothyriales* sp. KhNk32 str. CBS 129052, Vasse *et al*., 2017). Other abundant OTUs in these samples are classified as uncultured *Acetobacteraceae* (OTUs 26, 63, and 6325), Acidobacteria (*Bryocella*, OTU 104, and *Granulicella*, OTU 156), Actinobacteria (*Jartrophihabitans*, OTU 174, *Amnibacterium*, OTU 71, and *Curtobacterium*, OTU 7059), *Endobacter* (OTU 83), and *Sulfuriferula* (OTU 111). DO-1 and DO-2 cluster more closely with the IB and HB weathered rock samples than DO-3 and DO-4 (Fig. 2), and share a mixture of the OTUs abundant in both clusters.

### 2.3 Comparison to laboratory and field leaching experiments

We compared the microbial communities sampled here to rRNA gene libraries from weathering experiments described by Jones et al. (2017). These authors examined the microbial communities present in Duluth Complex waste rock and tailings from long-running laboratory and field leaching experiments (Lapakko and Antonson, 1994; Lapakko, 2015), including amplicon libraries from samples of laboratory humidity cell experiments (referred to here as “Lab” samples) and field weathering piles of simulated waste rock from pyrrhotite-containing Duluth Complex (“Field DC”) and pyrite-containing Ely Greenstone (“Field GS”) sources. For this comparison, we refer to the libraries from naturally weathered rock, seep sediment, and algal biomass for the present study as “outcrop” libraries to clearly separate them from libraries from the laboratory and field leaching experiments from the earlier work (Jones, Lapakko, *et al*., 2017). Information on all the libraries used in this comparison are summarized in Table S2.

In non-metric multidimensional scaling (NMS) analyses, libraries from outcrop samples (this study) separate from laboratory and field experiments (from Jones, Lapakko, *et al*., 2017) along the first ordination axis (Figure 3). Outcrop libraries are more similar to those from field experiments, with libraries from laboratory experiments plotting to the extreme of the first axis (Figure 3a). Some outcrop libraries overlapped with libraries from weathered Ely Greenstone. When laboratory experiments were excluded (Figure 3b), libraries from the Duluth Complex field leaching experiment still separate from the outcrop samples along the first ordination axis, but outcrop libraries also separate along both ordination axes, with libraries from the green seep mat (teal dotted circle) and seep sediment (blue solid circle) overlapping with the libraries generated from the Ely Greenstone field weathering experiments (purple open circles). Like in the hierarchical agglomerative cluster analysis (Figure 2), libraries from samples DO-3 and DO-4 (points enclosed by the black dashed line) cluster separately from the other outcrop samples as well as the laboratory and field experiments. Removal of OTUs classified as chloroplasts, as well as all chloroplast, mitochondria, and other eukaryote OTUs, did not affect the overall structure of the ordination (Figure S5).

Many OTUs co-occur in both the field weathering experiments and naturally-weathered outcrop samples. However, their abundances differ substantially. The OTUs that make up the most abundant communities in the naturally-weathered outcrop samples are present only at low abundances in the field weathering experiments. Accordingly, the OTUs that are most abundant in the field weathering experiments are present only at low abundances in the outcrop samples (Supp. Fig. S6).

### 2.4 Laboratory incubations

Because of the ubiquity of algae in the weathered outcrop samples, laboratory incubations were used to evaluate the effect of photosynthetic microorganisms on sulfide mineral weathering. Under acidic conditions, algae and heterotrophic organisms have been shown to diminish pyrite oxidation (see e.g., Johnson *et al*., 2008; Ňancucheo and Johnson, 2011), so incubations with isolate and enrichment cultures were conducted with and without algae to assess whether a similar decrease in pyrrhotite oxidation occurs under circumneutral conditions and how the presence of algae affects the bacterial community. Crushed pyrrhotite (sample Po2, Hobart *et al*., 2021) was incubated for 19 days with either an isolate or a mixed microbial community, with and without an algal inoculum. The isolate used in these experiments was *Sulfuriferula* sp. strain AH1, an autotrophic sulfur-oxidizing proteobacterium isolated from humidity cell experiments (Jones, Roepke, *et al*., 2017). The enrichment was a microbial community from weathered Duluth Complex rock samples IB-5 and GG-1, and mixed with a microbial community from lab and field experiments (Jones, Lapakko, *et al*., 2017). The algae inoculum was a mixture of material from samples IB-4, GG-2, and algae from a sulfate-reducing bioreactor described in Anderson (2018). The following treatments were performed, each in triplicate: (1) isolate only, (2) environmental inoculum only, (3) isolate plus algae, (4) environmental inoculum plus algae, and (5) algae alone. Samples with algae were incubated in the light on a diel cycle, and samples without were covered. At the termination of the experiment, 16S rRNA gene amplicon libraries were generated from the solids in order to evaluate which organisms were enriched, and how the presence of algae affected the community.

Libraries generated from the enrichment incubations were dominated by OTUs classified as members of the *Leeiaciae* family, with smaller amounts of *Sulfuriferula* spp., *Thiomonas* spp., and *Oxalicibacterium* spp. (Figure 4a). At the end of the experiment, all incubations inoculated with the algal biomass were dominated by OTUs classified as *Thiobacillus* (68-90%; Figure 4a), regardless of whether they started with the isolate, enrichment, or algal inoculum alone. OTUs identified as *Nitrospira, Nitrosomonas*, and *Sediminibacterium* were also abundant in these experiments. The algae inoculum was not a pure culture of algae strains but also included lithotrophic organisms, of which *Thiobacillus* was initially a minor component (0.04%) and was evidently enriched in the incubations (Figure 4a).

**Figure 4.**
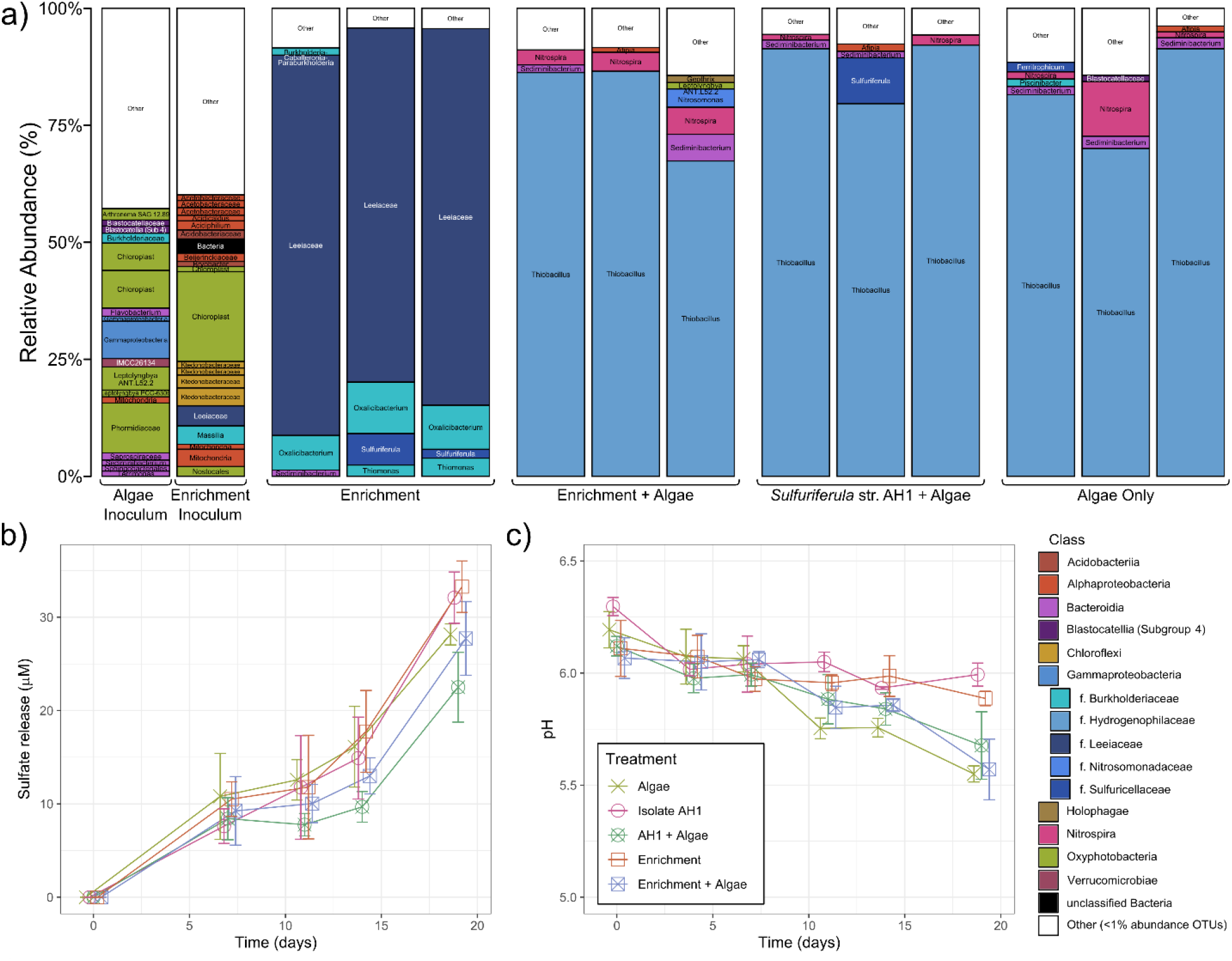
(a) Stacked bar chart showing taxonomic assignments of OTUs >1% relative abundance for the incubation experiments. “Other” represents those OTUs <1% abundance in each sample. Bars are colored by OTU taxonomic identification at the class level, with the exception of the *Gammaproteobacteria*, which are colored at the family level in varying shades of blue. Each bar is labeled with the OTU taxonomic classification at the genus level, or, at the highest available taxonomic classification. (b) Average sulfate release and (c) pH over the 19-day lifetime of the incubation experiments. Error bars represent standard deviations in sulfate concentrations or pH measured for the three treatment replicates. The x-axis position was shifted slightly in these plots to allow visualization of overlapping points.

Sulfate release was observed in all experiments, with slightly more sulfate release in the experiments without algae. By the end of the experiment, 18% less sulfate was released from environmental enrichments with algae than without, and 33% less sulfate was released from incubations with isolate AH1 with algae than without; however these differences are not statistically significant (Welch’s t-test, p > 0.05). In each incubation, pH dropped slightly over the course of the experiment, with those inoculated with algae measuring approximately 0.3 pH units lower than the incubations without algae at the termination. Only small concentrations of aqueous iron (II) were released over the course of the experiment (Fig. S7), with the AH1 + algae incubation showing the most variability in Fe^2+^.

## 3. DISCUSSION

### 3.1 Weathered rock-associated microbial communities

The microbial communities present on naturally-weathered outcrops of Duluth Complex material contain a diverse population of photosynthesizers, organotrophs, and lithotrophs. All libraries, even those collected from samples of seemingly bare weathered rock, contain substantial photosynthetic populations, with 6 OTUs representing chloroplasts among the top 30 most abundant OTUs. The DO samples, which were collected in early January, contain fewer OTUs classified as chloroplasts compared to the IB and HB samples that were collected in late June.

Microbial communities from the seep sediment (IB-2) and green algal mats (IB-4, GG-2) were distinct from weathered rock samples collected at the same site (IB-5, IB-6, GG-1). These libraries contain OTUs classified as the microaerophilic iron oxidizers *Leptothrix* and *Gallionella*, indicating that iron oxidation is a possible source of metabolic energy in the circumneutral seep environment. This is consistent with the reddish color of the seep sediment, which is indicative of the presence of iron oxides.

The microbial community from water-filled drill holes in consolidated rock at the gravel extraction site (sample GG-2) were dominated by OTUs representing euglenid algal chloroplasts and associated symbiotic organisms. *Euglena* spp. can be tolerant of high aqueous concentrations of metals and low pH, and are frequently found in AMD and other contaminated water systems (Brake *et al*., 2001; Casiot *et al*., 2004; Valente and Gomes, 2007; Das *et al*., 2009; Freitas *et al*., 2011; Jones *et al*., 2015). Their dominance in this sample, and their absence in other samples, is consistent with the higher metal content of waters that are in direct contact with the sulfide-rich rocks at site GG.

Microbial communities from the naturally-weathered outcrops and associated seeps are distinct from the communities that developed in laboratory and field weathering experiments described in earlier work (Jones, Lapakko, *et al*., 2017). The outcrop samples were most similar to experimentally-weathered Ely Greenstone (Figure 3). The most abundant sulfide in the Ely Greenstone is pyrite, while the sulfide mineral assemblage in the Duluth Complex is dominated by pyrrhotite and other acid soluble sulfides. However, the experimentally-weathered Duluth Complex field experiment had an average leachate pH of 4.7, while the pH of the Ely Greenstone field rock pile ranged from 7.2-7.3 (Jones, Lapakko, *et al*., 2017), closer to the pH 6.42 water measured in the effluent from the seep at the INCO site. This suggests that pH rather than the specific sulfide mineral assemblage is more important for determining microbial community composition in this setting. Further, we suspect that the distance between humidity cell experiments and the field piles and outcrop samples in multivariate statistical analyses (“Lab” in Fig. 3) is due to differences in organic material input and the corresponding effect on nitrogen and carbon cycling. The “Lab” humidity cell experiments were performed (by design) in an indoor environment with no or little low organic input, while both the field experiments and the naturally-weathered outcrops were from outdoor environments that would have had much higher organic input. Humidity cell experiments are designed to evaluate acid generation potential and mimic the weathering that would occur in the well-drained surface of a waste rock or tailings pile (Lapakko, 1988, 2015; Lapakko and Antonson, 2012), rather than weathering of reclaimed mine waste or in a natural forested setting.

Members of the class *Ktedonobacter* are abundant in the naturally-weathered outcrop samples and in the weathering field experiments (Field DC and Field GS in Figure 3 and S6). While numerous *Ktedonobacter* OTUs are present in either the outcrop samples or the field weathering experiments, three OTUs (23, 38, and 1091) are similar across all samples (Figure S4). OTU 1091 is most closely related (99.2% similarity) to an uncultured bacterium identified as part of a population of organisms that are early colonizers of volcanic rock deposits in Hawaii (Gomez-Alvarez *et al*., 2007), OTU 23 is most closely related (97.2% similarity) to a bacterial sequence from Antarctic soil (NCBI accession FR749799), and OTU 38 is most closely related (97.6% similarity) to an organism whose 16S rRNA gene was found in a chromium(IV) contaminated soil in China (NCBI accession KT016028). *Ktedonobacteria* appear to be ubiquitous in terrestrial environments, especially in organic-poor environments and during the early stages of soil formation (Delmont *et al*., 2015). The type strain for the genus *Ktedonobacter, Kt. racemifer* (SOSP1-21T), is an aerobic, filamentous, Gram-positive heterotroph isolated from the soil of a black locust wood in northern Italy (Chang *et al*., 2011). Organisms classified as *Ktedonobacter* have been identified in metagenome-assembled genomes in northern California grasslands (Butterfield *et al*., 2016), in Antarctic desert soils (Ortiz *et al*., 2021; Mezzasoma *et al*., 2022), and in hydrothermal sediments at an active deep-sea black smoker in the South Atlantic Ocean (Zhou *et al*., 2020). Related organisms in the same class have been isolated from a soil-like granular mass (Tengu-no-mugimeshi) collected from an alpine area in Gunma Prefecture, Japan (Wang *et al*., 2019), an ant nest collected in Honduras, soil from a pine wood in Spain, and soil “collected under a bush” in Corsica, France (Cavaletti *et al*., 2006). Yabe et al. (2021) describe the metabolic capabilities of seven *Ktedonobacter* isolates, all of which are mesophiles, heterotrophic, and form aerial mycelia, with an optimal growth pH of 5-7. Other members of this class, genus *Thermogemmatispora*, have been isolated from geothermally heated environments, and can oxidize carbon monoxide in addition to a diversity of carbon sources (Yabe *et al*., 2011; King and King, 2014). In the weathered rock environment sampled here, the *Ktedonobacteria* OTUs likely occupy a similar niche as part of the heterotrophic microbial community that forms early in the process of soil formation.

There were two OTUs identified as *Sulfuriferula* spp., a genus of sulfur-oxidizing Proteobacteria in the family *Sulfuricellaceae* that was abundant in weathered Duluth Complex rock from humidity cells and field experiments (Jones, Lapakko, *et al*., 2017). A *Sulfuriferula* strain isolated from the humidity cell experiments is an obligate autotroph that oxidizes inorganic sulfur compounds (Jones, Roepke, *et al*., 2017). The *Sulfuriferula* OTU that is most abundant in libraries from the field experiments (“Field DC”, Fig. S6) (OTU 8) makes up only a small fraction (less than 0.1%) of the OTUs found in the naturally-weathered “outcrop” libraries, while the *Sulfuriferula* OTU that is most abundant in libraries from the naturally-weathered outcrops (OTU 111) is less than 1% of the Field DC libraries (Figure S6). The genus *Sulfuriferula* contains multiple species that have different growth requirements, including both obligate autotrophs and mixotrophs, some of which can grow both aerobically and anaerobically (Watanabe *et al*., 2015, 2016; Jones, Roepke, *et al*., 2017; Kojima *et al*., 2020). Because of the diversity of metabolic capabilities in this genus, it would not be surprising if certain *Sulfuriferula* spp. have an advantage in the natural weathering environment, while different *Sulfuriferula* spp. have are more competitive in the lower organic input humidity cell experiments.

### 3.2 Impact of algae on microbial communities and sulfide weathering

Algae and other photosynthetic organisms have been explored as a tool for mitigating microbial breakdown of reactive sulfide minerals in a process broadly known as “bioshrouding” (Rawlings and Johnson, 2007; Johnson *et al*., 2008; Johnson, 2014; Bwapwa *et al*., 2017). Sulfide mineral oxidation by iron and sulfur-oxidizing bacteria can be mitigated by growing algae on the surface of reactive sulfide-rich tailings, because certain algae excrete organic compounds that are toxic to acidophilic autotrophs and also form an organic surface layer that limits oxygen penetration (Johnson *et al*., 2008; Johnson, 2014). Furthermore, the addition of algae and other photosynthetic organisms can have additional benefits for mitigating the impacts of acidic rock drainage: algae can accumulate and adsorb heavy metals, increase the alkalinity of the system, and produce organic acids such as lactic and acetic acid that mitigate the growth of acid-tolerant lithoautotrophs while providing food for heterotrophs (Peppas *et al*., 2000; Rambabu *et al*., 2020). However, these bioshrouding experiments have been performed on highly acidic materials, where mineral-oxidizing autotrophs are especially sensitive to the presence of organic acids that act as decouplers (Baker-Austin and Dopson, 2007). Here, we saw that incubation experiments with pyrrhotite at circumneutral pH did have slightly less sulfate release (and presumably less sulfide mineral dissolution) when a diverse algal culture was added to the experiments. However, the effects were relatively minor, as substantial sulfate release still occurred in the amended incubations.

One of the major effects of algal growth on the lithotrophic microbial community was the absence of *Sulfuriferula* OTUs and the dominance of *Thiobacillus* OTUs in the algae-amended incubations. The initial enrichment inoculum lacked OTUs identified as *Thiobacillus*, while the algae inoculum had 0.04% *Thiobacillus*. Conditions in the algae-amended incubations are evidently more favorable to *Thiobacillus*, perhaps because *Sulfuriferula* is more sensitive to organic compounds. Although the differences between sulfate release in each incubation were small, there was less sulfate released in the incubations inoculated with the *Sulfuriferula* strain AH1 and algae than in the experiment inoculated with *Sulfuriferula* alone, suggesting that the presence of algae and the algae-associated lithotrophs were able to outcompete the *Sulfuriferula* strain and result in less sulfide mineral dissolution. Future work will explore whether algal growth can be applied to limit sulfide mineral oxidation in mildly acidic or circumneutral waste like that anticipated from mining activities in the Duluth Complex.

### 3.3 Implications for weathering of Duluth Complex materials and other mixed sulfide ores

This study shows that the microbial communities that develop on naturally weathered Duluth Complex surfaces and in reclaimed rocks are distinct from those in leaching experiments (Figure 3). Microbial communities are sensitive indicators of environmental conditions, and so community composition can be used to compare how preexisting microbial communities change with varying levels of human disturbance. In this case, the microbial communities that are present before mining occurs could be used as a baseline to evaluate the successful reclamation of mine sites by the re-establishment of similar communities. The clear distinction between experimental and outcrop communities we observed here highlights the difference between the natural microbial communities and those that thrive in a disturbed environment. Conversely, the similarities between the microbial communities at the IB, GG, HB, and DO sites (Fig. 3a) suggests that environmental conditions at the historically disturbed IB and GG sites have converged with the more natural conditions at the HB and DO sites, and may be one indication of the effectiveness of the reclamation process at IB. However, more extensive sampling would be needed to fully characterize the existing microbial communities associated with the diverse ecosystems that have developed on natural and disturbed Duluth Complex material.

## ACKNOWLEDGEMENTS

The authors acknowledge Dean M. Peterson for field sampling assistance and insightful advice, and Satoshi Ishi for algae culturing advice and samples for use in the incubation experiments. Access to the IB and GG sampling sites was provided by the United States Forest Service and Twin Metals Minnesota. KKH was supported by a MnDRIVE Environment grant to JMF and DSJ, funding from the University of Minnesota Department of Earth & Environmental Sciences, and a Geological Society of America Student Research Grant. ZG and KH were supported by a NSF Research Experience for Undergraduates on Sustainable Land and Water Resources, coordinated by Diana Dalbotten.

The University of Minnesota is built on the ancestral lands of the Wahpekute band that was ceded to the United States by the Treaty of Traverse des Sioux in July of 1851, in an agreement that was not paid in full and whose underlying aim was the dissolution of the Dakota culture. The University has also benefited from Chippewa and Dakota (Mede-wakanton, Wahpekuta, Wahpeton and Sisseton Bands) land ceded by treaty and given to the University of Minnesota via the Morrill Act. Due to its land-grant status, the infrastructure, financial foundations, and faculty, students, and staff at the University of Minnesota all continue to benefit directly from these ceded lands, and we wish to acknowledge this support in our research.

