## Supplemental Information for "Microbial communities from weathered outcrops of a sulfide-rich ultramafic intrusion, and implications for mine waste management"

### MATERIALS AND METHODS

#### Field Samples

We collected weathered rock, sediment, and water samples from four locations in the South Kawishiwi Intrusion of the Duluth Complex (Severson, 1994): (i) roadside outcrops from the "Hammer Breaker" site along Forest Rte. 429 (HB samples, 47.7839849°, -91.792356°); (ii) a roadside outcrop off St. Louis Co. Road 623 (DO samples, 47.6892822°, -91.892129°); (iii) outcrops from a heavily weathered Duluth Complex surface that had been covered with glacial till until the 1940s (GG samples, 47.8311172°, -91.6779923°); and (iv) the reclaimed INCO pit (IB samples, 47.832392°, -91.67822°).

The "INCO Pit" site contains a small seep directly discharging from the reclaimed ore. Following reclamation in 1975, discharge was then intermittently monitored by the Minnesota

Environmental Quality Board and the U.S. Forest Service for water chemistry to evaluate specific risks from this site and understand the weathering of Duluth Complex material more broadly (e.g., Minnesota Environmental Quality Board, 1977). Seep discharge was sampled and analyzed for specific conductivity, alkalinity, turbidity, color, sulfate concentration, and dissolved cations (Fe, Ca, Mg, K, Na, Ni, Cu, Cd, Pb, Zn, Si) three times in the spring of 1976 and was found to have a pH of 6.8-7.0, with conductivities between 628-711 μS/cm and sulfate concentrations between 250 and 270 mg/L (see Minnesota Environmental Quality Board, 1977, for complete data). Similarly, pH, specific conductivity, alkalinity, and the concentration of aqueous metals (Fe, Ni, Cu, Cd, Pb, Zn) was measured in the nearby stream both upstream and downstream of the bulk sample site six times in the spring and summer of 1976. In those years, pH did not change significantly between the two sample sites (the upstream site had an average pH of 5.8, while the downstream site had an average pH of 5.7), and a statistically significant increase in aqueous copper and nickel concentration was found downstream of the bulk sample site (Minnesota Environmental Quality Board, 1977). Water from the site was sampled again in 2010 by the U.S. Forest Service, which found that the seep made a negligible contribution to sulfate and metal loading of nearby Filson Creek (Butcher, 2010).

We collected weathered rock from all sites, and seep sediment and water samples from sites IB and GG. Sample sites are summarized in Table 1 and Figure 1 in the main text, and in Figure S1 and S2, below. Field sampling was performed on 6 January 2016 (DO samples) and 28 June 2018 (IB, GG, and HB samples). Weathered rock and sediment samples were collected using sterile spatulas into sterile Whirl-Pac sample bags and sterile 50mL centrifuge tubes. For all sites, samples for DNA extraction were immediately frozen on dry ice after collection, and subsequently stored at -80°C until analysis. Samples for water chemistry were filtered through a 0.2μm polyethersulfone (PES) filter, and either preserved with 4M HCl and frozen or immediately frozen. Water samples for sulfide concentration were filtered into 15mL tubes containing pre-weighed 4mL aliquots of 5% w/v zinc chloride. No dissolved sulfide was detectable using the Cline assay using Hach reagents (Cline, 1969) (#2244500, Hach, Loveland, CO, USA).

**DNA extraction and amplicon library preparation.**

Total DNA from field samples and laboratory experiments was extracted using the PowerSoil DNA isolation kit (Qiagen, Hilden, Germany). To reduce DNA extraction bias, the vortexing step was modified, with aliquots removed after vortexing for 5, 10, and 15 minutes, and then recombined. Libraries were then prepared following the “in house” method of Jones, Lapakko et al. (2017). Briefly, the V4 region of the 16S rRNA gene was first amplified with primers “515f modified” and “806r modified” (Walters et al. 2015) that were amended with Nextera adaptors to allow barcoding (Jones, Lapakko, et al., 2017). PCR was performed as in Jones, Lapakko, et al. (2017): 5 min initial denaturation at 94°C, either 25 or 30 cycles of 45 s denaturation at 94°C, 60 s annealing at 50°C, and 90 s elongation at 72°C, and final elongation at 72°C for 10 min. Blank controls were included with all DNA extractions, and no product was visible in the blanks. PCR products were then submitted to the University of Minnesota Genomics Center for barcoding (10 cycles after 1:100 dilution) and sequencing on an Illumina MiSeq (Illumina, San Diego, CA, USA), 250 paired end cycles.

Raw libraries are available in the Sequence Read Archive at the National Center for Biotechnology Information (https://www.ncbi.nlm.nih.gov/sra) under accession PRJNA885421.

**Bioinformatic analyses.** OTU calling was performed as in Jones et al. (2021). Raw sequences were filtered and trimmed with Sickle (https://github.com/najoshi/sickle) to average quality above 28 (5' trimming only) and ≥ 100 bp; any residual adapters reverse complemented on the 3' end were removed with cutadapt (Martin, 2011); R1 and R2 reads were assembled with PEAR (Zhang et al., 2014); and primers removed by trimming the assembled reads with prinseq v.0.20.4 (Schmieder & Edwards, 2011). OTUs were defined at 97% similarity with a modified version of the UPARSE pipeline (USEARCH v.10.0; (Edgar, 2013)), in which the "derep_fulllength" script from VSEARCH v.1.9.5 (Rognes et al., 2016) was used. OTUs were classified with mothur v.1.36.1 (Schloss, 2020) to the Silva database v.132 (Quast et al., 2013) with a confidence score cutoff of 50.

Libraries were organized into a matrix of samples versus OTUs. Raw counts (i.e., the number of sequences of each OTU per sample) were first converted to proportional values to account for uneven library sizes by dividing by the total number of sequences in each library. OTUs that occurred at less than 0.01% were removed from the dataset. Prior to statistical analyses, the data were transformed using an arcsine square root transformation (Jones, Lapakko, et al., 2017). NMS ordinations were performed using 3 dimensions, with rotation to principal components, using the metaMDS() function in Vegan package v2.5-7 (Dixon, 2003) in R v4.1.0 (R Core Team, 2021) in RStudio (RStudio Team, 2021). Hierarchical agglomerative cluster analyses were performed with Bray-Curtis dissimilarity and unweighted pair-group method using arithmetic averages (UPGMA) clustering. Q-mode cluster analyses (clustering of samples) were calculated using all OTUS in the transformed dataset, while R-mode cluster analysis (clustering of OTUs) only included the top 30 most abundant OTUs. Plots were created using ggplot2 (Wickham, 2016).

**Laboratory Experiments.** Laboratory weathering experiments to test the effect of algal growth were conducted with crushed pyrrhotite as a substrate. Pyrrhotite used in these experiments was a mixture of 4C and 6C pyrrhotite with trace sphalerite, chalcopyrite, and galena from Ward's Scientific, crushed to a grain size of between 75 and 150µm. This corresponds to sample Po2 from (Hobart et al., 2021); see that work for a more complete description.

Five experimental treatments were conducted in triplicate in 50mL glass serum bottles: enrichment, enrichment + algae, isolate, isolate + algae, and algae only. 50mL of sterile growth media was added to each bottle, containing: 6mM $NH_4Cl$, 3mM $KH_2PO_4$, 1mM $Na_2HPO_4$, 1.5mM $MgCl_2 \cdot 6H_2O$, 0.3mM $CaCl_2 \cdot 2H_2O$, trace element solution (Flood et al., 2015), and 20mM MES buffer titrated to pH 6.0 with 4M NaOH. The isolate organism used in these experiments is *Sulfuriferula* sp. strain AH1 (Jones, Roepke, et al., 2017), an autotrophic sulfur-oxidizing β-proteobacteria isolated from humidity cell experiments described in Jones, Lapakko, et al. (2017). The enrichment inoculum was a homogenized microbial community collected from weathered Duluth Complex rock and previous incubation experiments, maintained in the laboratory in mixed culture with solid, crushed pyrrhotite as the growth substrate. The microbial communities identified through 16S rRNA libraries at the termination of the 19-day experiment are presented in Figure 4a in the main text, and sulfate release (as a proxy for sulfide mineral dissolution) is reported in Figure 4b. Experiments were placed on an orbital oscillating shaker exposed to a grow light on a 12 hour diel cycle. Those without the algae inoculum were covered to ensure continuous darkness.

Experiments were sampled every four to five days. At each sampling point, 6mL of leachate removed for chemical analysis and then replaced with 6mL of sterile growth media to maintain constant volume. 0.5mL of the leachate was analyzed for pH a LAQUAtwin pH-22 handheld pH meter (HORIBA, Kyoto, Japan); 1.5mL of the leachate was acidified with 20µm of 4M HCl and stored at -4℃ for ferrous iron analysis; 2mL of the leachate was stored at -20℃ for measurement of anion concentrations. Ferrous iron concentration was measured using the ferrozine assay (Stookey, 1970) using the method described in (Viollier et al., 2000). Anion concentrations (chloride, fluoride, bromide, nitrate, sulfate) were measured using a Metrohm 930 Compact IC Flex ion chromatograph with a A Supp 5 column, 20µL sample loop, and an eluent carbonate buffer (3.2mM $Na_2CO_3$ and 1.0mM $NaHCO_3$). Sulfate release was calculated based on the measured concentration of sulfate at each time point and subsequently correcting for the leachate replaced with sulfate-free media at each sampling interval. Statistical significance was assessed with Welch's t-test using the R function t.test().

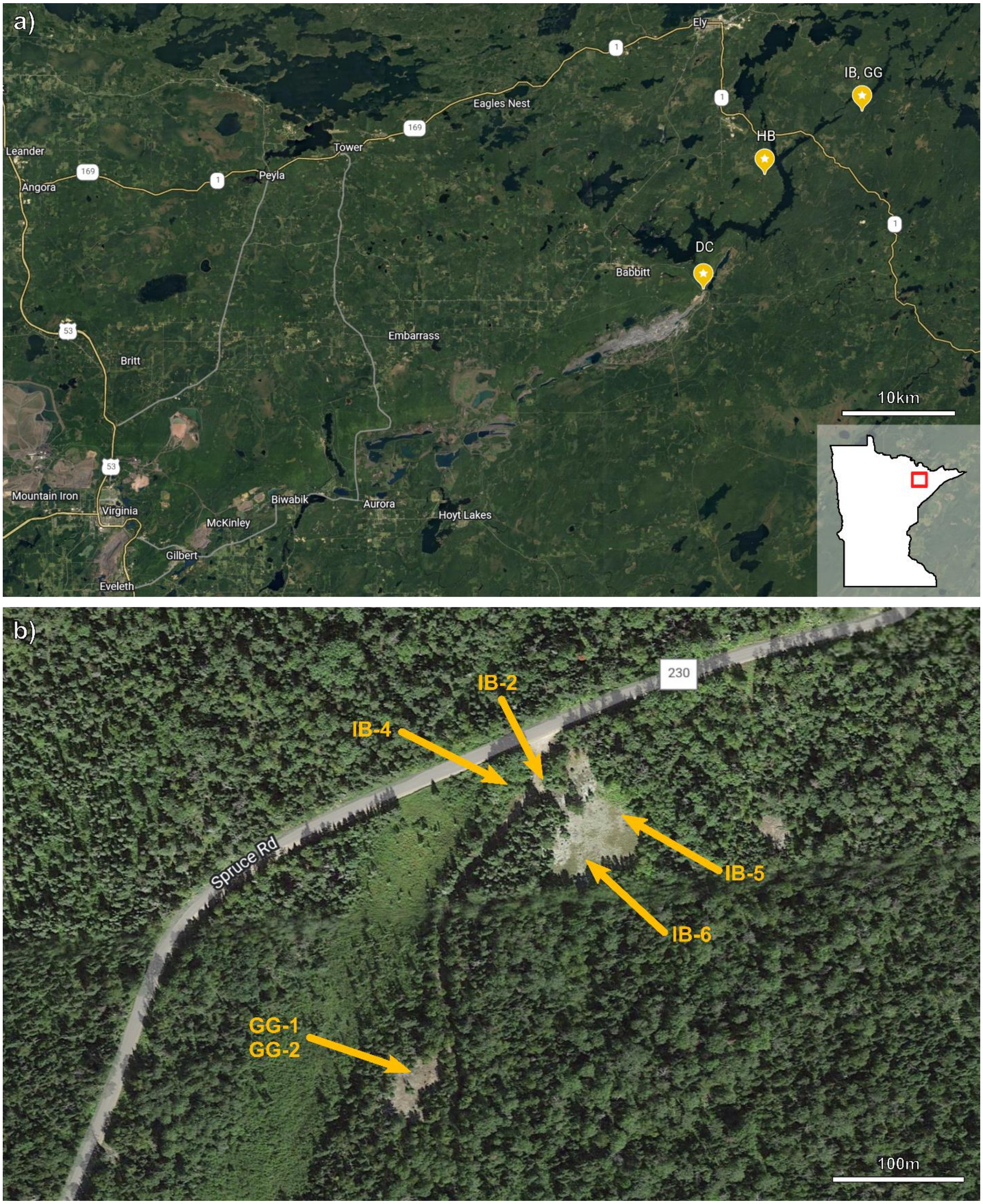


**Figure S1.** (a) Map of sampling site locations. (b) Sampling locations at the IB/GG site. Map data from Google Earth; imagery from LANDSAT/Copernicus (NOAA).

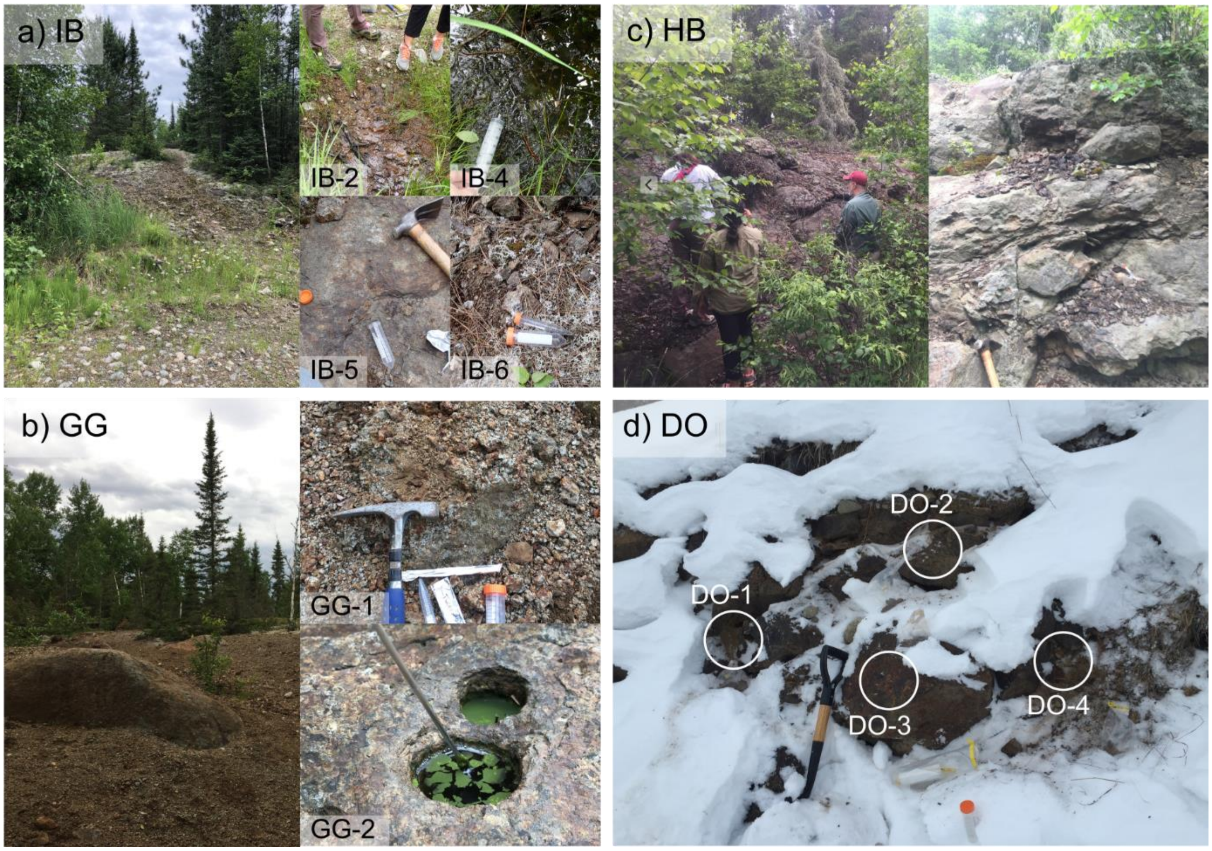

**Figure S2.** Sample site images for (a) IB, (b) GG, (c) HB, and (d) DO sites.

**Table S1.** Concentration of dissolved anions measured by IC for aqueous samples collected at the IB and GG sampling sites. bd = below detection limit. No detectable phosphate, bromide, or nitrite.

| | Sample Type | Fluoride (µM) | Chloride (µM) | Nitrate (µM) | Sulfate (µM) |
|---|---|---|---|---|---|
| **IB-2** | Effluent from seep | 52.4 | 729.4 | 65.3 | 477.3 |
| **IB-4** | Water from stream below seep | 31.0 | 304.4 | bd | 581.6 |
| **GG-2** | Water from drill hole in consolidated DC rock | 31.0 | 305.9 | bd | 360.0 |

**Table S2.** 16S rRNA gene libraries from Jones, Lapakko, et al. (2017) re-analyzed alongside the libraries from this study.

| Sample ID | Formation | Experiment type | Material | % S | Category (this study) |
|---|---|---|---|---|---|
| DCW-HC9a | Duluth Complex | Humidity cell | Crushed rock | 0.13 | Lab |
| DCW-HC13a | Duluth Complex | Humidity cell | Crushed rock | 0.55 | Lab |
| DCW-HC13a | Duluth Complex | Humidity cell | Crushed rock | 0.55 | Lab |
| DCW-HC15a | Duluth Complex | Humidity cell | Crushed rock | 1.03 | Lab |
| DCW-HC15a | Duluth Complex | Humidity cell | Crushed rock | 1.03 | Lab |
| DCW-HC17a | Duluth Complex | Humidity cell | Crushed rock | 0.23 | Lab |
| DCW-HC19a | Duluth Complex | Humidity cell | Crushed rock | 0.61 | Lab |
| DT-HC8a | Duluth Complex | Humidity cell | Tailings | 0.2 | Lab |
| DT-HC8a | Duluth Complex | Humidity cell | Tailings | 0.2 | Lab |
| HF15-2 | Duluth Complex | Field rock pile | Crushed rock | 0.9 | Field DC |
| HF15-2 | Duluth Complex | Field rock pile | Crushed rock | 0.9 | Field DC |
| HF15-3 | Duluth Complex | Field rock pile | Crushed rock | 0.9 | Field DC |
| HF15-4 | Duluth Complex | Field rock pile | Crushed rock | 0.9 | Field DC |
| HF15-5 | Duluth Complex | Field rock pile | Crushed rock | 0.9 | Field DC |
| HF15-7 | Duluth Complex | Field rock pile | Crushed rock | 0.9 | Field DC |
| HF15-8 | Duluth Complex | Field rock pile | Crushed rock | 0.9 | Field DC |
| HF15-9 | Duluth Complex | Field rock pile | Crushed rock | 0.9 | Field DC |
| HF15-10 | Duluth Complex | Field rock pile | Crushed rock | 0.9 | Field DC |
| HF15-10 | Duluth Complex | Field rock pile | Crushed rock | 0.9 | Field DC |
| HF15-11 | Duluth Complex | Field rock pile | Crushed rock | 0.9 | Field DC |
| HF15-12 | Duluth Complex | Field rock pile | Crushed rock | 0.9 | Field DC |
| HF15-14 | Duluth Complex | Field rock pile | Crushed rock | 0.9 | Field DC |
| HF15-15 | Duluth Complex | Field rock pile | Crushed rock | 0.9 | Field DC |
| GS-HC1a | Ely Greenstone | Humidity cell | Crushed rock | 0.04 | Lab |
| GS-HC4a | Ely Greenstone | Humidity cell | Crushed rock | 0.1 | Lab |
| GS-HC6a | Ely Greenstone | Humidity cell | Crushed rock | 0.12 | Lab |
| GS-HC8a | Ely Greenstone | Humidity cell | Crushed rock | 0.16 | Lab |
| GS-HC10a | Ely Greenstone | Humidity cell | Crushed rock | 0.2 | Lab |
| GS-HC18a | Ely Greenstone | Humidity cell | Crushed rock | 1.22 | Lab |
| GS-HC18a | Ely Greenstone | Humidity cell | Crushed rock | 1.22 | Lab |
| HF15-18 | Ely Greenstone | Field rock pile | Crushed rock | | Field GS |
| HF15-19 | Ely Greenstone | Field rock pile | Crushed rock | | Field GS |
| HF15-20 | Ely Greenstone | Field rock pile | Crushed rock | | Field GS |
| HF15-21 | Ely Greenstone | Field rock pile | Crushed rock | | Field GS |
| HF15-23 | Ely Greenstone | Field rock pile | Crushed rock | | Field GS |
| HF15-23 | Ely Greenstone | Field rock pile | Crushed rock | | Field GS |
| HF15-25 | Ely Greenstone | Field rock pile | Crushed rock | | Field GS |
| HF15-26 | Ely Greenstone | Field rock pile | Crushed rock | | Field GS |

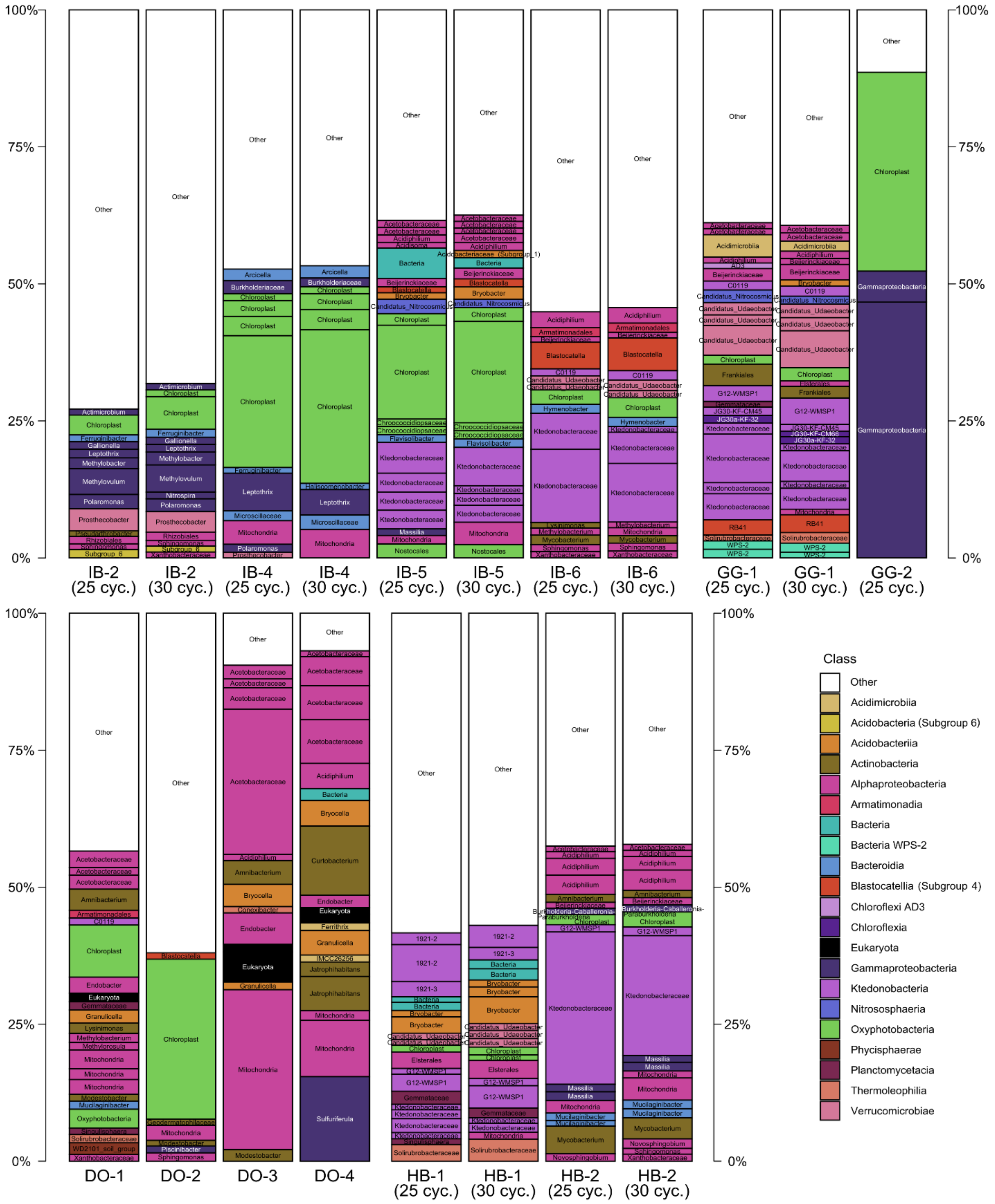


**Figure S3.** OTU abundances >1% for the IB, GG, DO, and HB amplicon libraries. OTUs are identified at the genus level, or, at the highest available taxonomic level. OTUs are colored by their classification at the class (or highest available) level. Columns labeled “25 cyc.” are the libraries generated from the sample after 25 PCR cycles, and columns labeled “30 cyc.” are the libraries generated from the sample after 30 PCR cycles.

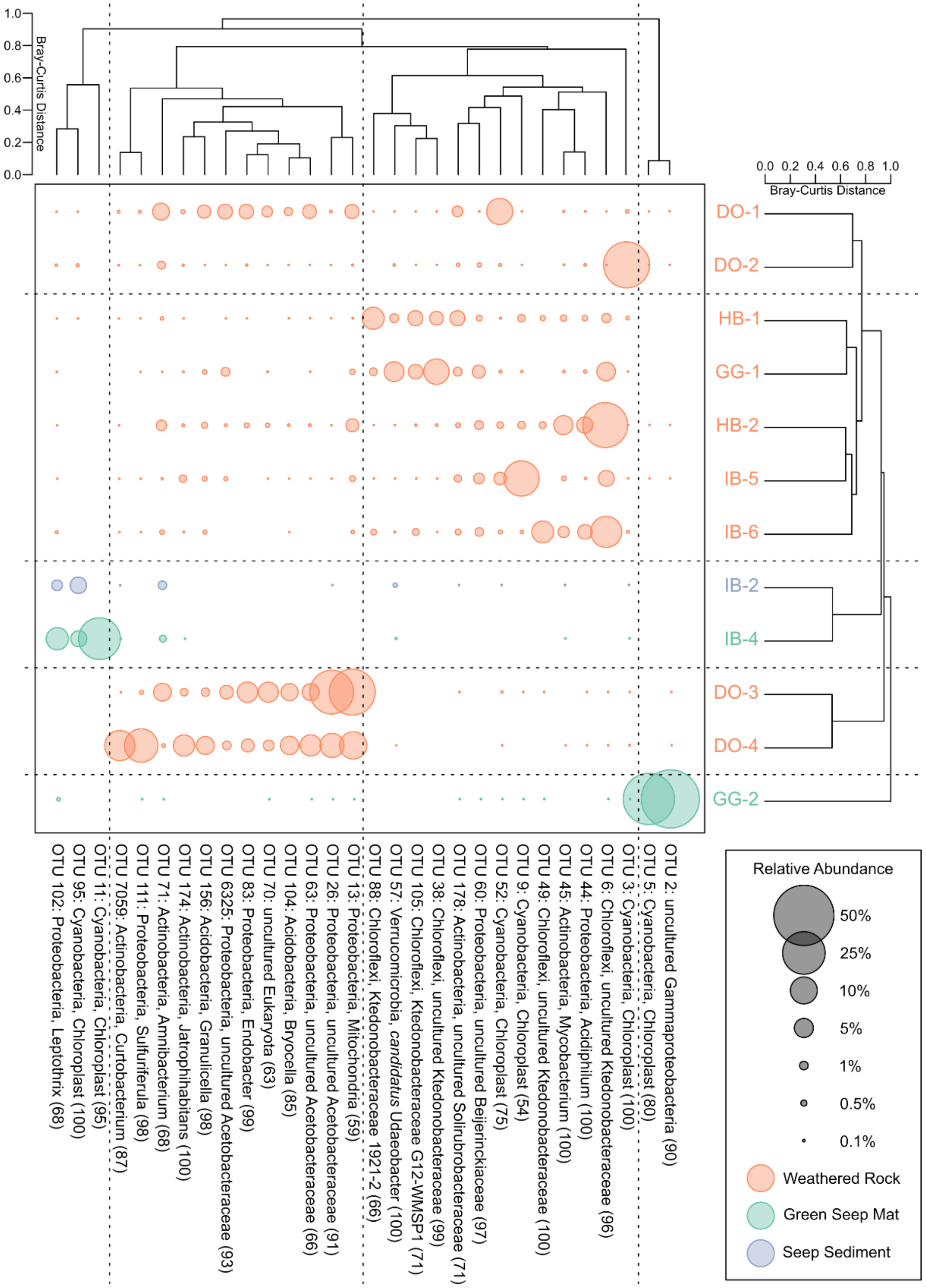


**Figure S4.** Hierarchical agglomerative cluster analysis of libraries collected for this study. Size of the points scales with the relative abundance of the OTUs. The Q-mode cluster analysis was calculated with all OTUs, while the R mode cluster analysis only included the top 30 most abundant OTUs. The taxonomic affiliation of each OTU includes its phylum- and genus-level classification, with confidence scores provided in parentheses. OTUs that are unclassified at the genus level are identified with the highest available taxonomic classification.

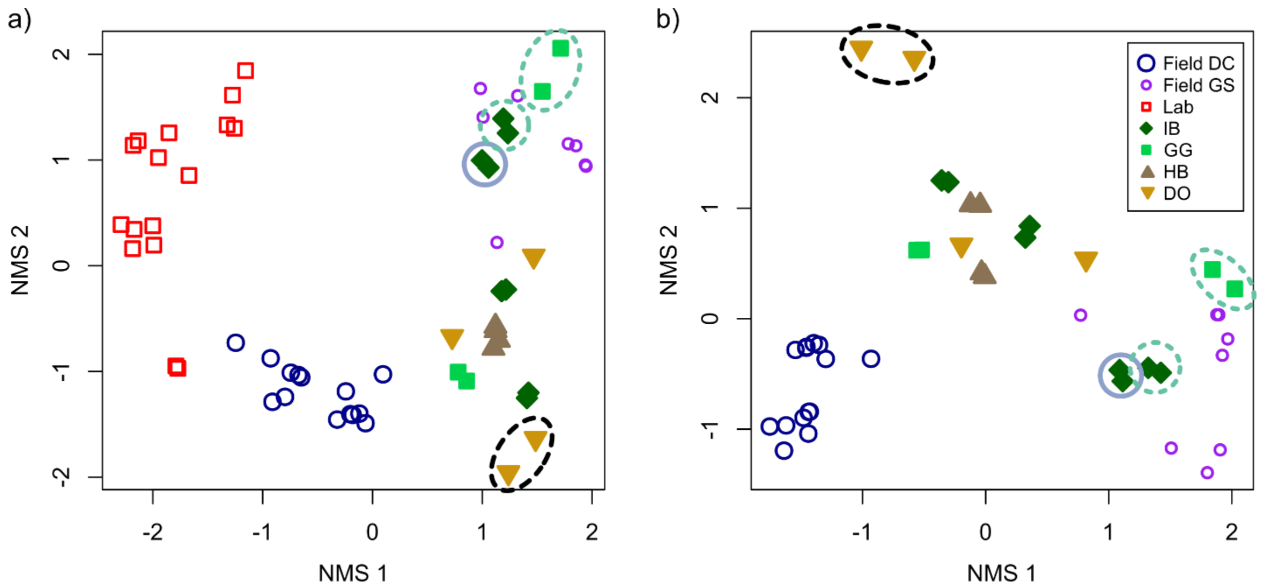


**Figure S5.** (a) NMS ordinations of rRNA amplicon libraries from naturally-weathered Duluth Complex outcrops (libraries IB, GG, HB, and DO, this study), laboratory humidity cells ("Lab", red open circles), and experimental field rock piles. This analysis includes the chloroplast, mitochondria, and Eukaryote OTUs removed in Fig. 3. The experimental field rock piles include field leaching experiments with Duluth Complex material ("Field DC," dark blue larger circles) and Ely Greenstone ("Field GS," purple smaller circles). The "Lab" and "Field" leaching experiments are from Jones et al. (2017a). (b) NMS ordinations of rRNA amplicon libraries from the experimental field rock piles and naturally weathered outcrops only, with all OTUs classified as chloroplasts removed. Stress for (a) is 7.5, stress for (b) is 5.8. Sky blue solid circled points indicate seep sediment samples, teal green dotted circled points indicate water and algal biomass samples. All other IB, GG, HB, and DO samples were collected from weathered rock surfaces. The black dashed circle indicates DO samples that are separated from the other weathered rock samples in the hierarchical agglomerative cluster analysis (Figure 2).

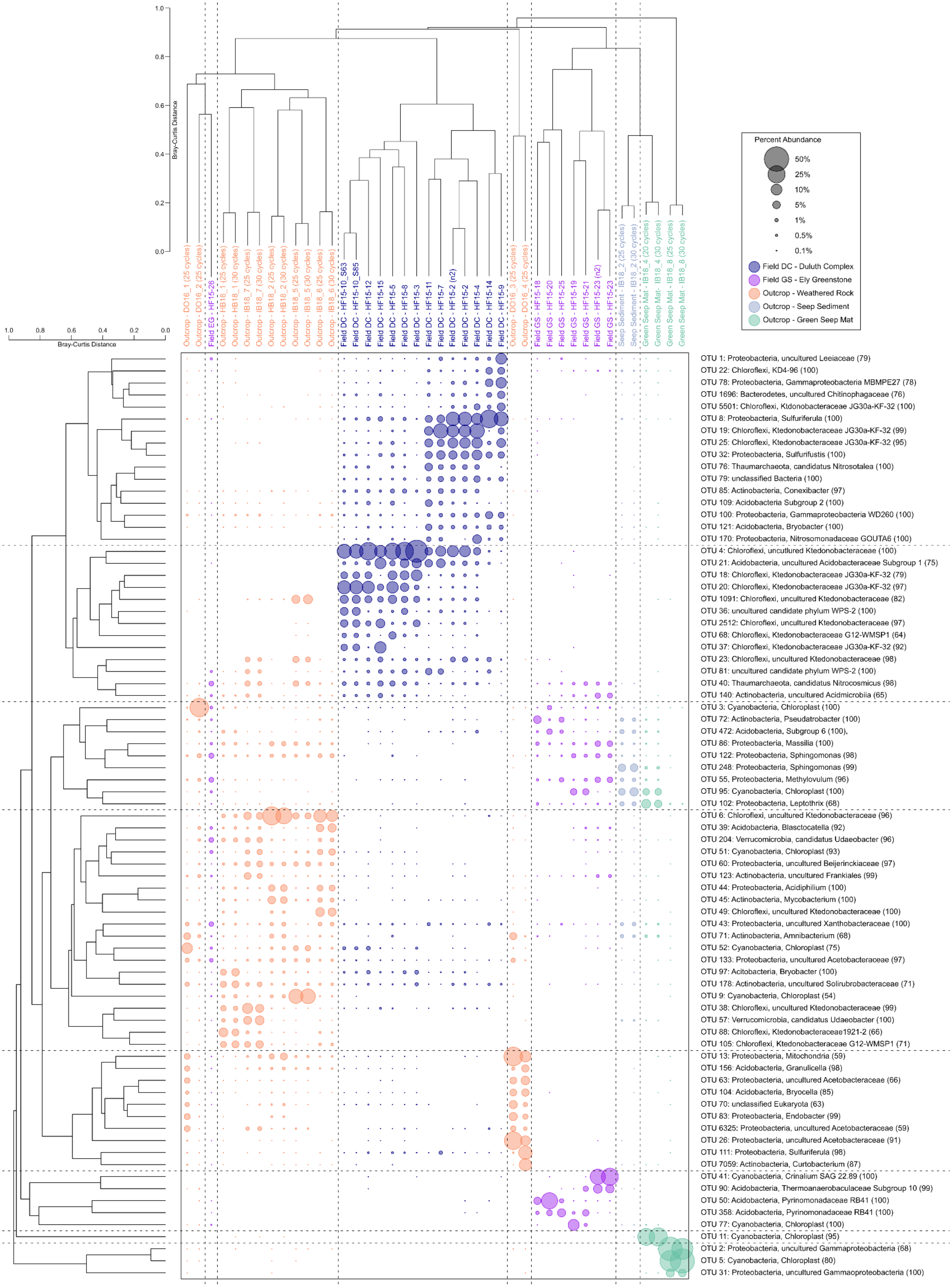

Bray-Curtis Distance
1.0
0.8
0.6
0.4
0.2
0.0
Percent Abundance
50%
25%
10%
5%
1%
0.5%
0.1%
Field DC - Duluth Complex
Field GS - Ely Greenstone
Outcrop - Weathered Rock
Outcrop - Seep Sediment
Outcrop - Green Seep Mat
Outcrop - DO16_1 (25 cycles)
Outcrop - DO16_2 (25 cycles)
Field EG - HF15-26
Outcrop - HB18_1 (25 cycles)
Outcrop - HB18_1 (30 cycles)
Outcrop - IB18_7 (25 cycles)
Outcrop - IB18_7 (30 cycles)
Outcrop - HB18_2 (25 cycles)
Outcrop - HB18_2 (30 cycles)
Outcrop - IB18_5 (25 cycles)
Outcrop - IB18_5 (30 cycles)
Outcrop - IB18_6 (25 cycles)
Outcrop - IB18_6 (30 cycles)
Field DC - HF15-10_S63
Field DC - HF15-10_S85
Field DC - HF15-12
Field DC - HF15-15
Field DC - HF15-5
Field DC - HF15-8
Field DC - HF15-3
Field DC - HF15-11
Field DC - HF15-7
Field DC - HF15-2 (n2)
Field DC - HF15-2
Field DC - HF15-4
Field DC - HF15-14
Field DC - HF15-9
Outcrop - DO16_3 (25 cycles)
Outcrop - DO16_4 (25 cycles)
Field GS - HF15-18
Field GS - HF15-20
Field GS - HF15-25
Field GS - HF15-19
Field GS - HF15-21
Field GS - HF15-23 (n2)
Field GS - HF15-23
Seep Sediment - IB18_2 (25 cycles)
Seep Sediment - IB18_2 (30 cycles)
Green Seep Mat - IB18_4 (20 cycles)
Green Seep Mat - IB18_4 (30 cycles)
Green Seep Mat - IB18_8 (25 cycles)
Green Seep Mat - IB18_8 (30 cycles)
OTU 1: Proteobacteria, uncultured Leeiaceae (79)
OTU 22: Chloroflexi, KD4-96 (100)
OTU 78: Proteobacteria, Gammaproteobacteria MBMPE27 (78)
OTU 1696: Bacterodetes, uncultured Chitinophagaceae (76)
OTU 5501: Chloroflexi, Ktdonobacteraceae JG30a-KF-32 (100)
OTU 8: Proteobacteria, Sulfuriferula (100)
OTU 19: Chloroflexi, Ktedonobacteraceae JG30a-KF-32 (99)
OTU 25: Chloroflexi, Ktedonobacteraceae JG30a-KF-32 (95)
OTU 32: Proteobacteria, Sulfurifustis (100)
OTU 76: Thaumarchaeota, candidatus Nitrosotalea (100)
OTU 79: unclassified Bacteria (100)
OTU 85: Actinobacteria, Conexibacter (97)
OTU 109: Acidobacteria Subgroup 2 (100)
OTU 100: Proteobacteria, Gammaproteobacteria WD260 (100)
OTU 121: Acidobacteria, Bryobacter (100)
OTU 170: Proteobacteria, Nitrosomonadaceae GOUTA6 (100)
OTU 4: Chloroflexi, uncultured Ktedonobacteraceae (100)
OTU 21: Acidobacteria, uncultured Acidobacteraceae Subgroup 1 (75)
OTU 18: Chloroflexi, Ktedonobacteraceae JG30a-KF-32 (79)
OTU 20: Chloroflexi, Ktedonobacteraceae JG30a-KF-32 (97)
OTU 1091: Chloroflexi, uncultured Ktedonobacteraceae (82)
OTU 36: uncultured candidate phylum WPS-2 (100)
OTU 2512: Chloroflexi, uncultured Ktedonobacteraceae (97)
OTU 68: Chloroflexi, Ktedonobacteraceae G12-WMSP1 (64)
OTU 37: Chloroflexi, Ktedonobacteraceae JG30a-KF-32 (92)
OTU 23: Chloroflexi, uncultured Ktedonobacteraceae (98)
OTU 81: uncultured candidate phylum WPS-2 (100)
OTU 40: Thaumarchaeota, candidatus Nitrocosmicus (98)
OTU 140: Actinobacteria, uncultured Acidmicrobiia (65)
OTU 3: Cyanobacteria, Chloroplast (100)
OTU 72: Actinobacteria, Pseudatrobacter (100)
OTU 472: Acidobacteria, Subgroup 6 (100),
OTU 86: Proteobacteria, Massilia (100)
OTU 122: Proteobacteria, Sphingomonas (98)
OTU 248: Proteobacteria, Sphingomonas (99)
OTU 55, Proteobacteria, Methylovulum (96)
OTU 95: Cyanobacteria, Chloroplast (100)
OTU 102: Proteobacteria, Leptothrix (68)
OTU 6: Chloroflexi, uncultured Ktedonobacteraceae (96)
OTU 39: Acidobacteria, Blasctocatella (92)
OTU 204: Verrucomicrobia, candidatus Udaeobacter (96)
OTU 51: Cyanobacteria, Chloroplast (93)
OTU 60: Proteobacteria, uncultured Beijerinckiaceae (97)
OTU 123: Actinobacteria, uncultured Frankiales (99)
OTU 44: Proteobacteria, Acidiphilium (100)
OTU 45: Actinobacteria, Mycobacterium (100)
OTU 49: Chloroflexi, uncultured Ktedonobacteraceae (100)
OTU 43: Proteobacteria, uncultured Xanthobacteraceae (100)
OTU 71: Actinobacteria, Amnibacterium (68)
OTU 52: Cyanobacteria, Chloroplast (75)
OTU 133: Proteobacteria, uncultured Acetobacteraceae (97)
OTU 97: Acitobacteria, Bryobacter (100)
OTU 178: Actinobacteria, uncultured Solirubrobacteraceae (71)
OTU 9: Cyanobacteria, Chloroplast (54)
OTU 38: Chloroflexi, uncultured Ktedonobacteraceae (99)
OTU 57: Verrucomicrobia, candidatus Udaeobacter (100)
OTU 88: Chloroflexi, Ktedonobacteraceae1921-2 (66)
OTU 105: Chloroflexi, Ktedonobacteraceae G12-WMSP1 (71)
OTU 13: Proteobacteria, Mitochondria (59)
OTU 156: Acidobacteria, Granulicella (98)
OTU 63: Proteobacteria, uncultured Acetobacteraceae (66)
OTU 104: Acidobacteria, Bryocella (85)
OTU 70: unclassified Eukaryota (63)
OTU 83: Proteobacteria, Endobacter (99)
OTU 6325: Proteobacteria, uncultured Acetobacteraceae (59)
OTU 26: Proteobacteria, uncultured Acetobacteraceae (91)
OTU 111: Proteobacteria, Sulfuriferula (98)
OTU 7059: Actinobacteria, Curtobacterium (87)
OTU 41: Cyanobacteria, Crinalium SAG 22.89 (100)
OTU 90: Acidobacteria, Thermoanaerobaculaceae Subgroup 10 (99)
OTU 50: Acidobacteria, Pyrinomonadaceae RB41 (100)
OTU 358: Acidobacteria, Pyrinomonadaceae RB41 (100)
OTU 77: Cyanobacteria, Chloroplast (100)
OTU 11: Cyanobacteria, Chloroplast (95)
OTU 2: Proteobacteria, uncultured Gammaproteobacteria (68)
OTU 5: Cyanobacteria, Chloroplast (80)
OTU 31: Proteobacteria, uncultured Gammaoproteobacteria (100)

**Figure S6 (previous page).** Hierarchical agglomerative clustering analysis of “field” samples from Jones et al. (2017a) with “outcrop” samples from this study. The Q-mode cluster analysis was calculated with all OTUs, while the R-mode cluster analysis only included OTUs appearing at >10% in any one sample. The taxonomic affiliation of each OTU includes its phylum- and genus-level classification, with confidence scores provided in parentheses. OTUs that are unclassified at the genus level are identified with the highest available taxonomic classification. Outcrop samples (this study) collected from weathered rock are in orange, samples of water and green seep biomass are in teal green, and samples of seep sediments are in sky blue. Samples from experimental field piles of Duluth Complex material (Field DC) are in dark blue, and samples from field piles of Ely Greenstone material (Field EG) are in purple.

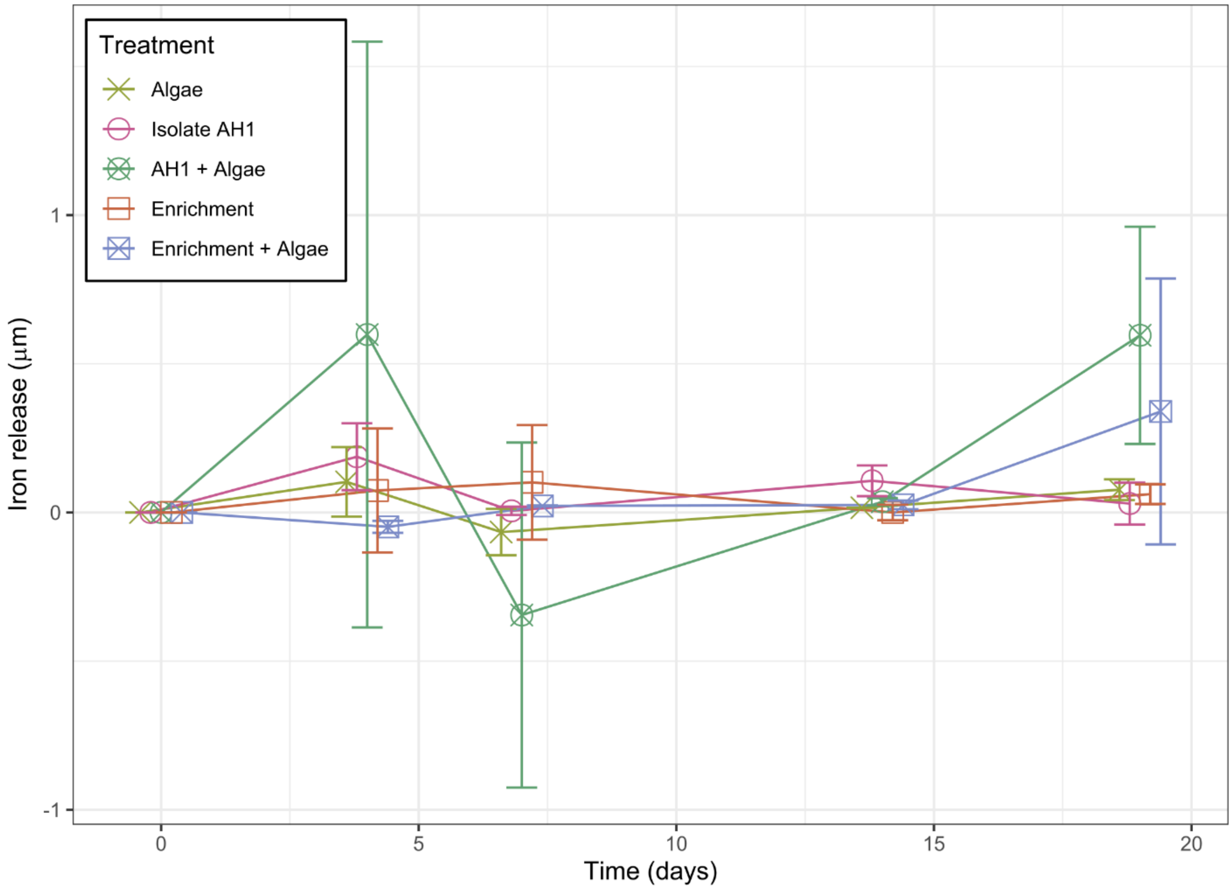


**Figure S7.** Average aqueous iron (II) release over the 19-day lifetime of the incubation experiments. Dissolved iron (II) concentration was measured colorimetrically by the ferrozine assay.
